# An AI-Driven Platform for Deconstructing and Engineering Biomolecular Recognition

**DOI:** 10.64898/2025.12.09.692808

**Authors:** Fodil Azzaz, Jacques Fantini

**Affiliations:** Aix-Marseille Univ, INSERM UA16, Marseille, France

## Abstract

Understanding molecular interactions at the atomic level remains a central challenge across biology, medicine, and engineering. We introduce Perturbation Scanning (PS), an interrogative AI framework that actively deconstructs molecular interfaces. PS integrates graph-based representations of structures derived from molecular dynamics trajectories or Protein Data Bank files to systematically probe the electrostatic, hydrophobic, and steric contributions of each residue. To translate these insights into actionable design, we introduce the Intelligent Interface Optimization Scanner (IIOS), a standalone tool that generates energy-scored mutation proposals from interface maps. Together, PS and IIOS provide an integrated platform for dissecting and rationally engineering molecular interactions by resolving force-specific and stage-dependent contributions that are not directly accessible with existing computational approaches. Unlike traditional alanine scanning or free-energy methods such as MMPBSA—which provide only static or ensemble-averaged measures—PS delivers stage-resolved, force-wise decomposition of binding interfaces via a suite of targeted physicochemical perturbations applied to its AI model, directly quantifying each residue’s mechanistic role.

## Introduction

Understanding the physicochemical principles that govern molecular recognition requires moving beyond structural observation toward systematic interrogation. Molecular dynamics simulations capture interaction dynamics(1,2) , and machine learning models predict binding affinities, but these approaches remain fundamentally observational: they describe what happens during binding without actively testing which residues drive recognition through specific physical forces.

The central challenge is the transition from correlation to mechanism. Structural snapshots(3) and affinity measurements can identify residues associated with function, while free-energy calculations(4) provide ensemble-averaged contributions. However, these methods lack the resolution to decompose interfaces into their underlying electrostatic, hydrophobic, and steric components or to track how these contributions evolve across binding stages.

This limitation has broad practical consequences. In drug discovery(5), the inability to distinguish essential from peripheral contacts hinders rational optimization. In synthetic biology(6), engineering novel protein interfaces remains difficult without a clear understanding of the physical determinants of recognition. Even in well-studied systems such as neurotoxin–receptor interactions(7,8), differential binding can be observed, yet its physical basis often remains unresolved.

Current computational approaches—including molecular dynamics(1,2), free-energy calculations(4), and machine learning predictors— either remain observational or provide aggregated measures of binding. While these methods successfully describe interaction patterns or predict affinities, they lack a unified framework to resolve how distinct physicochemical forces contribute to binding across different temporal stages. Similarly, although protein design tools exist for stability or affinity optimization, they do not directly translate mechanistic interface profiles into ranked, force-specific mutation proposals. Here we close these gaps with Perturbation Scanning (PS) and the Intelligent Interface Optimization Scanner (IIOS). PS overcomes these limitations by integrating a graph neural network with a systematic perturbation engine, enabling residue-wise, force-specific, and stage resolved attribution of binding mechanisms.

PS employs a trained graph model as a surrogate system, enabling counterfactual interrogation via targeted physicochemical perturbations (e.g., charge reversal, hydrophobicity inversion). These perturbations independently modify electrostatic, hydrophobic, and steric properties across defined binding stages, directly quantifying residue and force-specific contributions. The framework operates on both molecular dynamics trajectories and static structures (PS-Lite), extending its utility across diverse systems. To convert mechanistic insight into actionable design, we integrate PS with the Intelligent Interface Optimization Scanner (IIOS), which transforms interface profiles into ranked, energetically scored mutation proposals. Together, PS and IIOS establish a unified paradigm for the mechanistic understanding and rational engineering of biomolecular interfaces (Fig. 1).

**Fig 1.**
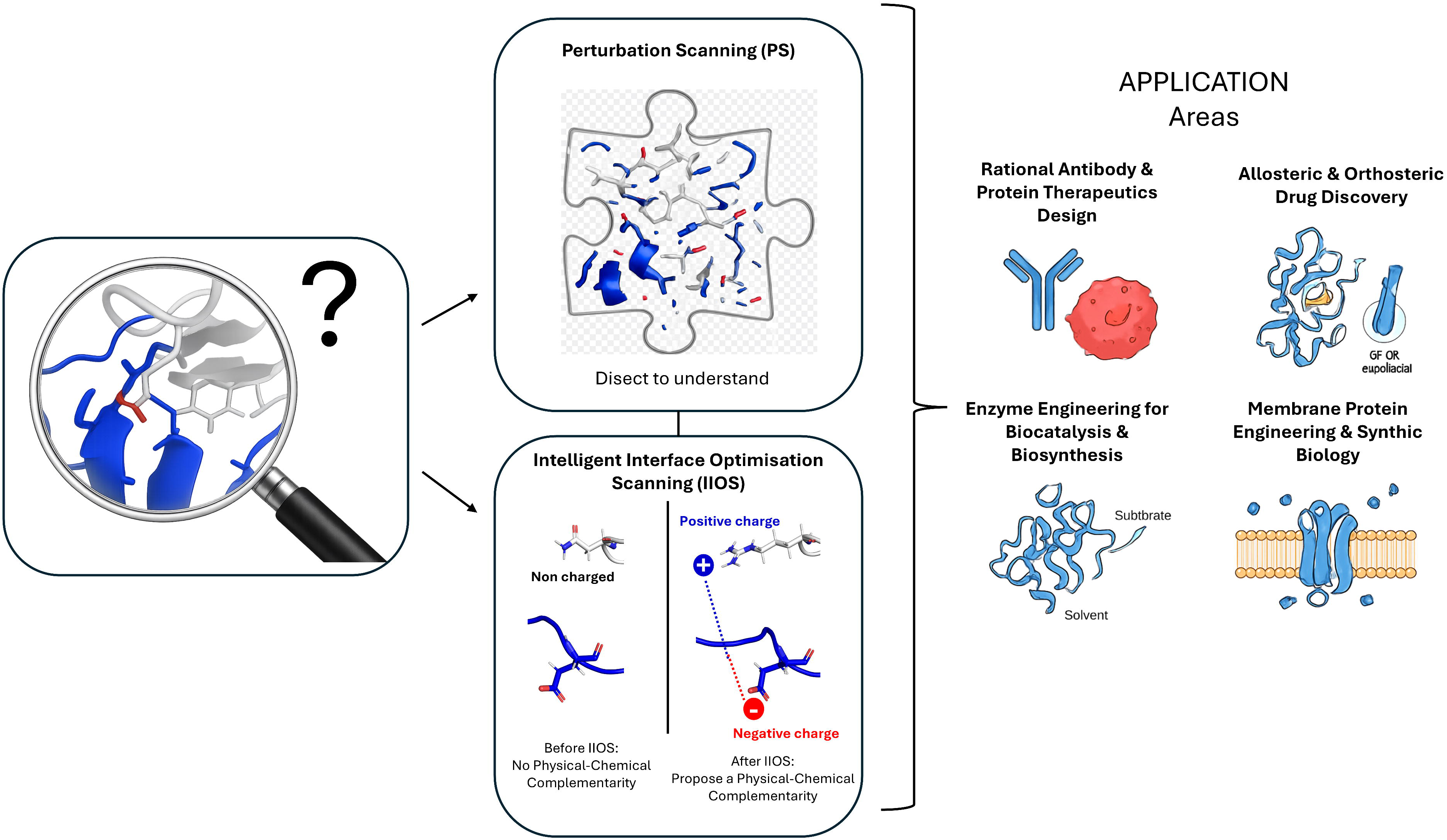
Overview of the Perturbation Scanning (PS) and Intelligent Interface Optimization Scanner (IIOS) framework. Left, a biomolecular interface is interrogated beyond structural observation, highlighting the central question of how specific residues contribute to recognition. Center, Perturbation Scanning (PS) deconstructs molecular interfaces by applying systematic, force-specific in silico perturbations to a graph-based AI surrogate model trained on molecular dynamics trajectories or static structures. Residue-wise perturbations (e.g., electrostatic, hydrophobic, steric) are applied across user-defined binding stages, enabling mechanistic attribution of how and when individual residues contribute to interface stability. Bottom center, Intelligent Interface Optimization Scanner (IIOS) translates PS-derived physicochemical interface maps into actionable design proposals by scanning interfacial and near-interface residues and suggesting energetically scored mutations that improve physical–chemical complementarity. Right, representative application areas include rational antibody and protein therapeutic design, allosteric and orthosteric drug discovery, enzyme engineering for biocatalysis and biosynthesis, and membrane protein engineering and synthetic biology. Together, PS and IIOS form an integrated platform that closes the loop from mechanistic interface deconstruction to rational biomolecular engineering.

## Results and Discussion

Rather than introducing new experimental datasets, we deliberately benchmark PS against multiple, independent, experimentally established binding phenomena across toxin–receptor and membrane systems, using agreement with known biological behavior as a stringent validation criterion.

### The PS Framework: An AI Engine for Mechanistic Interrogation

We developed Perturbation Scanning (PS), a graph-based AI framework that transforms molecular interaction analysis from passive observation to active mechanistic interrogation. PS represents biomolecular systems as graphs in which atoms are encoded with physicochemical feature vectors—including electrostatics, hydrophobicity, mass, and structural context (9,10)—and edges represent covalent bonds or non-covalent contacts (<6 Å), identified via k-d tree algorithms (11). These graphs are processed by a graph convolutional network (GCN) (12) trained with frame interleaving, using batch normalization and dropout for regularization (13), to predict interface stability. The model learns hierarchical representations that integrate covalent and non-covalent interactions across conformational ensembles (**Fig. S1**).

Robustness was validated through 10 independent training trials with bootstrap confidence intervals (95% CI), assessing R², Spearman correlation, mean absolute error, and within-tolerance accuracy. The final model was selected based on balanced validation loss and test R², ensuring predictive reliability.

### Configurable Temporal Staging Enables Stage-Resolved Mechanistic Deconstruction

A defining innovation of PS is its configurable temporal staging, which allows interaction analysis to be partitioned into user-defined binding stages. For clarity in this study, we divided trajectories into three equal-length regimes—early (initial binding events), mid (equilibrium interactions), and late (stabilized complex formation)—but the framework supports any number of stages, uneven time intervals, or unsupervised clustering to identify natural kinetic phases. This flexibility permits PS to model diverse dynamic processes, including stepwise complex assembly, induced-fit binding, and dissociation pathways, aligning the analysis with the underlying biochemical timeline.

### Interface Perturbation Scanning Quantifies Mechanistic Contributions

To move beyond correlation toward mechanistic attribution, PS implements six perturbation strategies: electrostatic charge reversal, hydrophobic inversion, bulky side-chain substitution, π-stacking disruption, hydrogen-bond network breaking, and structural displacement. For each residue, the perturbation effect Δ is quantified as the absolute change in predicted interface stability: Δ = |P(perturbed) – P(original)|. Residues are ranked by the sum of Δ values across perturbations (Sum Δ), weighting both the magnitude and the frequency of interaction perturbation. This approach directly attributes binding contributions to specific physicochemical forces, answering not only *which* residues matter but *how* and *when* they contribute(14).

### Computational Efficiency: Mechanistic Insight from a Single Trajectory

A key practical advantage of PS is its dramatic computational efficiency. Traditional methods for identifying critical residues typically require multiple independent MD simulations of mutant variants—each consuming hundreds of GPU hours and weeks of researcher time. For comprehensive analysis of interfaces like those studied here (20–30 residues across multiple binding stages), this translates to months of computational effort and substantial financial cost. In contrast, PS extracts comparable mechanistic insights relevant to residue-level and force-specific contributions through *in silico* perturbations applied to a single wild-type trajectory, avoiding the need for separate mutant simulations and reducing computational requirements by approximately N-fold (where N is the number of residues studied). This efficiency gain enables rapid comparative analysis across multiple systems—as demonstrated below—that would be computationally prohibitive through brute-force mutation simulations, democratizing detailed mechanistic analysis for laboratories without access to supercomputing resources.

### BoNT/B1–SYT1 and BoNT/B1–SYT2 Systems Reveal Distinct Assembly Architectures

Applied to the BoNT/B1–SYT1 and BoNT/B1–SYT2 neurotoxin–receptor systems, PS captured fundamentally different dynamic landscapes. For the flexible SYT1 complex, the model achieved a mean test R² of 0.81 ± 0.035 with Spearman ρ = 0.90 ± 0.013, reflecting broad conformational fluctuations. In contrast, the rigid SYT2 complex yielded R² ≈ 0 due to minimal variance across the trajectory, yet maintained substantial rank correlation (ρ = 0.704 ± 0.031), demonstrating PS’s ability to adapt to systems with varying degrees of conformational freedom (Fig. S2A–L).

Applying the PS perturbation suite revealed that SYT1 employs a GT1b-dependent compensatory assembly strategy (Fig. 4A–B). Initial SYT1–GT1b interactions weaken over time (Δ = 0.472 → 0.144), compensated by strengthening BoNT/B1–GT1b contacts (Δ = 0.647 → 1.123), indicating a dynamic handoff mechanism where the ganglioside transitions from SYT1 recruitment to BoNT/B1 stabilization (Fig. 2C). This GT1b-mediated handoff mechanistically explains the experimentally established requirement for ganglioside presence in BoNT/B1 binding to SYT1, revealing how this dependence operates dynamically during assembly (15). Key residues driving this compensation include LYS-1187/1188 and GLU-1189 at the BoNT/B1-SYT1 interface, and R1242 engaging GT1b’s sialic acid and N-acetylgalactosamine moieties (Fig. 2D).

**Fig 2.**
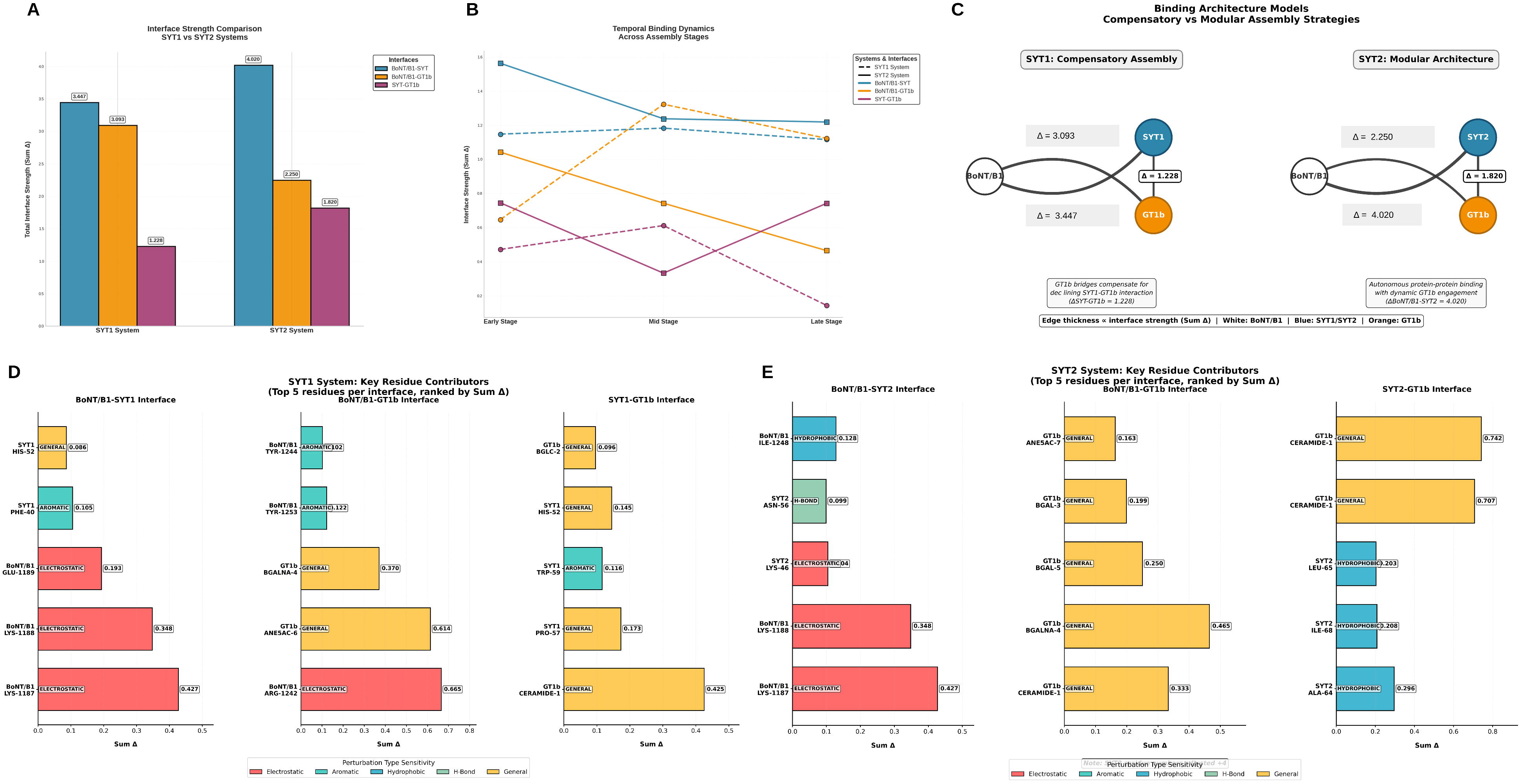
Comparative Interface for BoNT/B1-SYT1-GT1b and BoNT/B1-SYT2-GT1b systems. Total interface strength comparison showing SYT2’s stronger autonomous protein-protein binding (blue) versus SYT1’s compensatory GT1b bridging (orange) (A). Temporal dynamics reveal SYT1’s compensatory handoff (dashed lines) versus SYT2’s modular stability (solid lines) (B). Network architecture models illustrate compensatory (SYT1) versus modular (SYT2) assembly strategies; edge thickness ∝ interface strength (C). Top residue contributors for SYT1 and SYT2 systems, ranked by Sum Δ and colored by perturbation type sensitivity (D-E).

**Fig 3.**
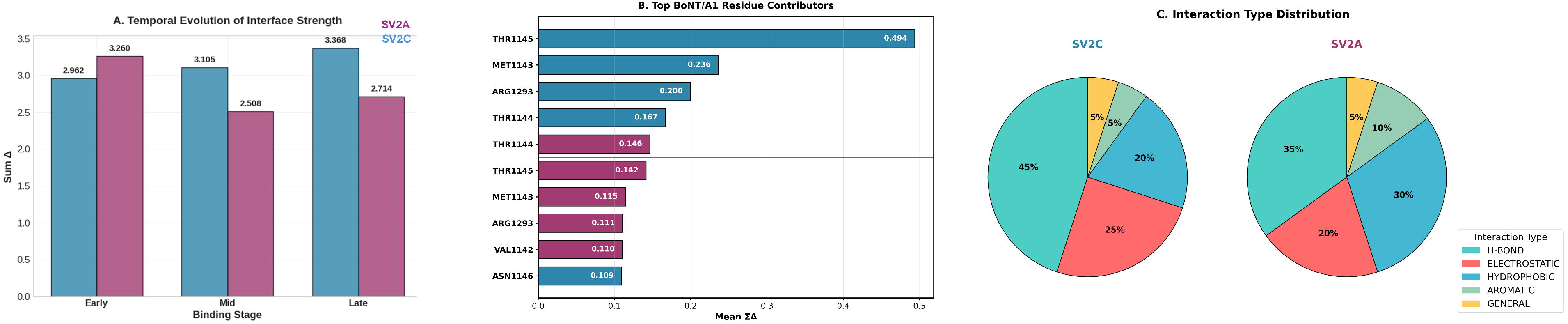
Perturbation Scanning Analysis of BoNT/A1 Binding to SV2A vs SV2C. Bar plots show the Sum Δ (interface perturbation sensitivity) for SV2C (blue) and SV2A (purple) complexes across early, mid and late binding stages (A). Horizontal bar chart siplaying the mean Sum Δ values for the top five contributing residues from BoNT/A1 in each complex (B). Pie charts comparing the percentage contribution of different physicochemical interaction types to total interface strength (C). Hsd is for histidine, BGALNA is for β-D-N-acetylgalactosamine, and BGAL is for β-galactose. The numbers reflect their position in the sequence (from 1 to 7).

**Fig 4:**
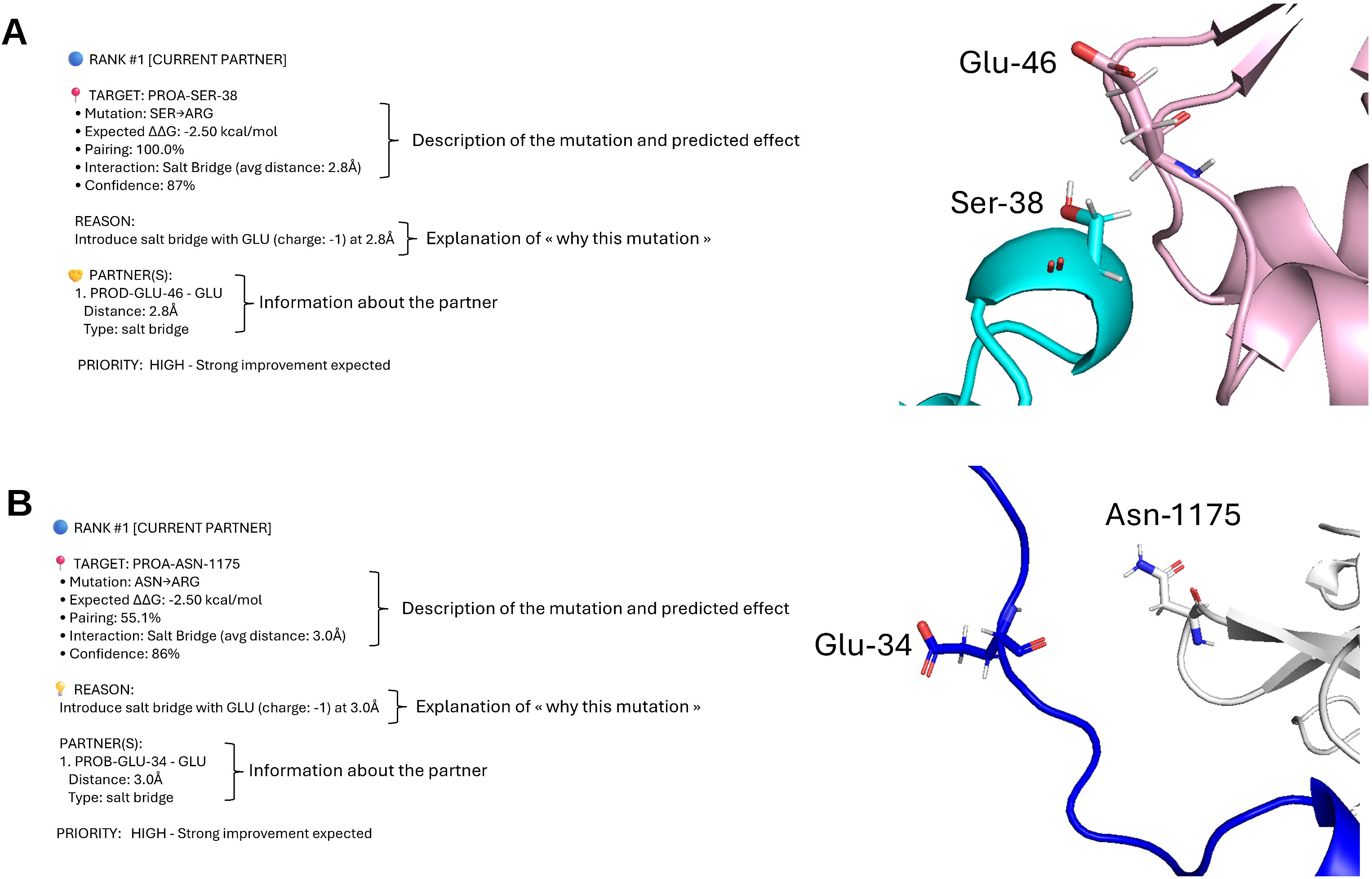
Exemple of raw output provided by Intelligent Interface Optimization Scanner (IIOS) for Barnase-Barstar (A) and BoNT/B1-SYT2 complex (B).

In contrast, SYT2 exhibits a modular, autonomous binding architecture, with a stronger BoNT/B1 interface (Sum Δ = 4.020) and a recoverable U-shaped SYT2–GT1b engagement. This autonomy is stabilized by K1187/1188, K46, and N56, with ganglioside contacts maintained through a hydrophobic patch (A64, L65, I68) (Fig. 2E).

Together, these findings illustrate how PS decodes not only interaction patterns but the physical mechanisms underlying differential receptor specificity. PS provides a mechanistic deconstruction of known experimental observations: SYT1’s ganglioside dependence emerges from a GT1b-bridged compensatory assembly, whereas SYT2 engages via a direct, protein-centric modular interface.

### BoNT/A1–SV2A and BoNT/A1–SV2C Systems Unveil Divergent Temporal Engagement Strategies

Expanding to BoNT/A1–SV2A and BoNT/A1–SV2C complexes, PS uncovered strikingly different binding dynamics. The SV2C interface demonstrated progressive stabilization, with perturbation sensitivity increasing from Δ = 2.962 (early) to 3.368 (late), indicating continuous optimization. Conversely, the SV2A complex exhibited transient instability, starting strong (Δ = 3.260 early), dropping sharply at mid-stage (2.508), then partially recovering (2.714 late), suggesting rearrangement or competitive displacement before weaker final contacts form (Fig. 3A).

Residue-level analysis highlighted Thr-1145 as a dominant contributor in SV2C (Δ = 0.4938) versus a much weaker role in SV2A (0.1416), with Met-1143 and Arg-1293 also showing enhanced contributions in the high-affinity complex (Fig. 3B). Interaction-type profiling revealed that SV2C binding is hydrogen-bond-dominated (45% of interface strength), while SV2A exhibits a more balanced distribution (35% H-bonds, 30% hydrophobic, 20% electrostatic) (Fig. 3C). These findings align with and mechanistically explain the experimentally established higher binding affinity of BoNT/A1 for SV2C over SV2A(16). PS reveals the biophysical mechanism behind SV2C’s superior affinity: through a progressively optimized hydrogen-bond network that strengthens across binding stages, while SV2A relies on transient, balanced interactions that fail to stabilize over time.

### Plasmolipin–Lipid Interactions: Lipid-Specific Resolution in Membrane Environments

Beyond protein–protein systems PS provides per-lipid resolution in analyzing membrane protein environments, enabling quantitative comparison of individual lipid contributions. Applied to plasmolipin embedded in a lipid-raft membrane (POPC/cholesterol/GM1) versus a pure POPC bilayer(17), PS revealed ∼4-fold higher interaction strengths in the raft system (early Δ = 33.99 vs. 8.37 in POPC) (Table S1). Strikingly, ceramide (CER160), despite constituting only 13.1% of contact frequency, contributed 35.1% of the total Δ score (16.48 Δ), demonstrating a disproportionate stabilizing role. In contrast, POPC (44.3% contact frequency) accounted for only 21.4% (10.06 Δ). Ganglioside polar components and cholesterol collectively contributed 39.3% of the total interaction score (Table S2). This lipid-resolved insight moves beyond phenomenological descriptions of “raft stabilization” to quantify how specific lipid chemistries orchestrate membrane protein stability through stage-dependent interactions, establishing a quantitative basis for understanding lipid selectivity and therapeutic targeting in membrane biology.

### PS-Lite: Accessible Interface Analysis for Static Structures

To broaden accessibility, we developed PS-Lite, a deterministic, non–machine-learning variant designed for rapid and interpretable analysis of static structures. PS-Lite applies the same physicochemical feature engineering as full PS but replaces the neural network with rule-based scoring based on contact density and residue properties. Validation on the Barnase–Barstar complex (PDB: 1BRS) correctly identified experimentally critical residues (Barnase Arg-59, Arg-87; Barstar Asp-35, Asp-39). Applied to BoNT/A1–SV2C (PDB: 6ES1), it highlighted key interface residues including SV2C Lys-558, Phe-563, and Asn-559, and BoNT/A1 Tyr-1122, Tyr-1149, and Thr-1145/1146. While lacking temporal resolution, PS-Lite provides rapid, interpretable interface mapping suitable for mutagenesis guidance, preliminary screening, and educational use, extending the framework’s utility to researchers without MD capabilities.

### IIOS Identifies Actionable Mutation Opportunities

To translate PS-derived interface insights into rational design, we applied the Intelligent Interface Optimization Scanner (IIOS). For the Barnase–Barstar complex, IIOS generated 48 prioritized mutation suggestions, including SER38→ARG (ΔΔG_pred = −2.5 kcal/mol) and ARG59→PHE (ΔΔG_pred = −2.3 kcal/mol) to establish new salt bridges and π-stacking interactions. Proximity scanning further identified under-utilized interfacial positions such as LEU89→ARG and THR100→PHE, highlighting the potential to create novel contacts (Fig. 4A).

Applied to a BoNT/B1–SYT2 molecular dynamics trajectory, IIOS proposed 34 mutations, with ASN1175→ARG and GLN1176→ARG predicted to strengthen electrostatic complementarity near SYT2 Glu-rich regions. Aromatic enhancements (e.g., THR1182→PHE) were suggested to reinforce π-stacking with SYT2 Phe-48, while proximity scanning revealed opportunities for new π-stacking interactions at near-interface residues such as ILE1259→PHE/TYR (Fig. 4B). These proposals illustrate how IIOS systematically converts structural and dynamic interface profiles into energetically scored design blueprints, bridging mechanistic understanding with practical protein engineering.

### PS as a Complementary Analytical Layer for Mechanistic Interface Analysis

The development of PS represents a shift from observational to interrogative computational structural biology. Unlike molecular dynamics simulations—which describe but do not explain—or free-energy methods that provide ensemble-averaged contributions, PS actively quantifies the role of individual residues through systematic *in silico* perturbations. Unlike machine-learning predictors that correlate structure with affinity, PS attributes binding contributions to specific electrostatic, hydrophobic, and steric forces across user-defined kinetic stages. This stage-resolved, force-wise deconstruction has no direct counterpart in current computational toolkits. By integrating graph-based representation, strategic frame interleaving, and multi-faceted *in silico* mutagenesis, PS moves beyond identifying important residues to deconstructing their stage-wise physicochemical contributions. Benchmarking across four neurotoxin–receptor systems spanning two toxin families (BoNT/B1 with α-helix recognition and BoNT/A1 with β-strand engagement) demonstrates PS’s generality and robustness. The framework successfully captured compensatory assembly (SYT1), modular autonomy (SYT2), progressive stabilization (SV2C), and transient instability (SV2A), revealing that molecular recognition is not merely a static complementarity but an orchestrated sequence of physical interactions.

### Biological Implications and Therapeutic Design

The differential binding strategies decoded by PS have direct implications for understanding botulism pathogenesis and developing targeted interventions. For example, SYT1’s ganglioside-dependent compensatory mechanism suggests that engineered soluble decoys could disrupt the handoff step, while SYT2’s protein-centric modular interface might be blocked by high-affinity peptide inhibitors. Similarly, SV2C’s progressive hydrogen-bond network provides a template for designing stable binders, whereas SV2A’s transient interface may require allosteric disruptors. More broadly, PS reveals that binding sequence—the temporal order of interaction formation—may be as important as binding affinity for biological specificity, aligning with emerging systems-biology perspectives on stage-coordinated function. PS does not claim experimental causality; rather, it provides a controlled counterfactual interrogation of physicochemical contributions that can guide and rationalize experimental design.

### Limitations and Future Directions

While PS represents a significant advance, several limitations warrant consideration. First, perturbations are computational; experimental validation of key predictions across multiple systems is needed. Second, analysis of individual trajectories could be strengthened by integrating multiple replicas for more robust statistics. Third, stage definition, though flexible, requires user input; automated clustering approaches could further streamline analysis. Fourth, future applications to systems with large conformational changes, disordered regions, or multi-domain rearrangements will further test the framework’s robustness. Directions for extension include: (1) incorporating experimental data via multi-modal learning, (2) developing adaptive stage-segmentation algorithms, (3) extending perturbation strategies to post-translational modifications and allosteric effects, and (4) establishing standardized benchmarks for stage-resolved interaction analysis across broader molecular classes. The modular architecture of PS ensures that these extensions can be implemented without fundamental redesign. PS is intended as a hypothesis-generating and mechanistic interpretation framework, not a replacement for experimental validation.

### Conclusion: From Observational Science to Engineering Discipline

Perturbation Scanning (PS) provides a generalizable, computationally efficient framework for deconstructing biomolecular interfaces into their fundamental physical components across user-definable binding stages. By revealing not only *which* residues participate but *how* and *when* they contribute through specific forces, PS transforms our capacity to decode the complex language of molecular recognition. In doing so, PS moves beyond the aggregated statistics of free-energy calculations and the static approximations of traditional mutagenesis tools, introducing a stage-aware analytical framework for mechanistic biomolecular engineering. Coupled with the standalone design tool IIOS, the pipeline closes the loop from mechanistic insight to rational interface engineering. As structural biology increasingly captures transient states and AI generates dynamic ensembles, frameworks like PS will be essential for extracting mechanistic understanding from complex datasets. By making stage-dependent mechanisms interpretable and computationally accessible, PS shifts molecular recognition from an observational science to an engineering discipline—enabling researchers not only to understand but to actively orchestrate biomolecular interactions across diverse biological contexts.

## Supporting information

Materiel and method, Figure S1-2, Table S1-2

## Author Contributions

F.A.: Conceptualization, Methodology, Software, Validation, Formal Analysis, Investigation, Data Curation, Writing – Original Draft, Writing – Review & Editing, Visualization.

J.F.: Supervision, Writing – Original Draft, Writing – Review & Editing, Resources.

## Data and Materials Availability

The Perturbation Scanning (PS) software suite, including PS-Lite and IIOS, is publicly available as well-documented, user-ready source code at https://github.com/fodil13.

The repository includes tutorials, example configurations, and centralized parameter files, enabling researchers to apply the framework to their own systems with minimal setup. The code is released under a custom open-source license that permits free academic use; commercial use requires a separate license. All data needed to evaluate the conclusions in the paper are present in the paper, the Supplementary Materials.

---

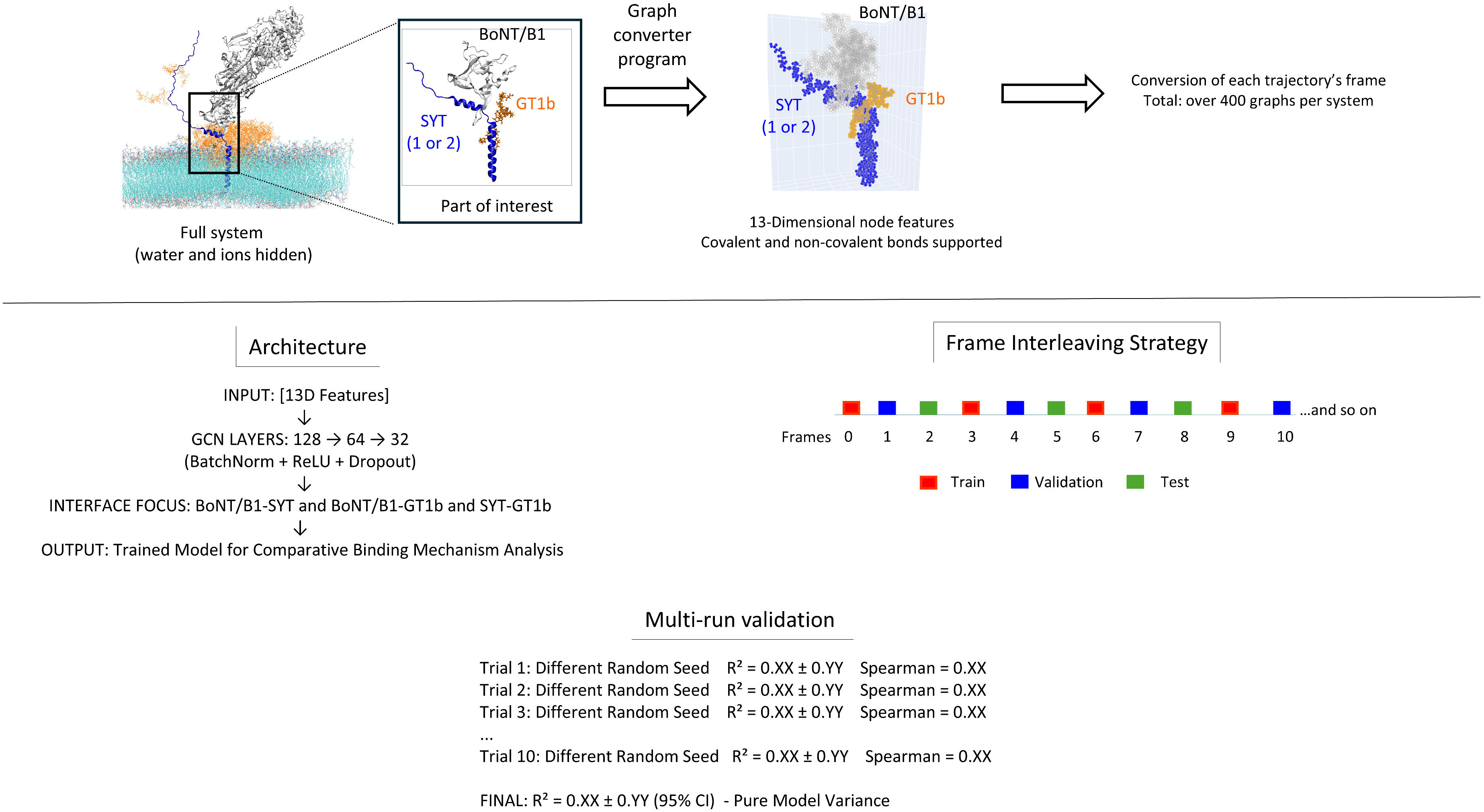

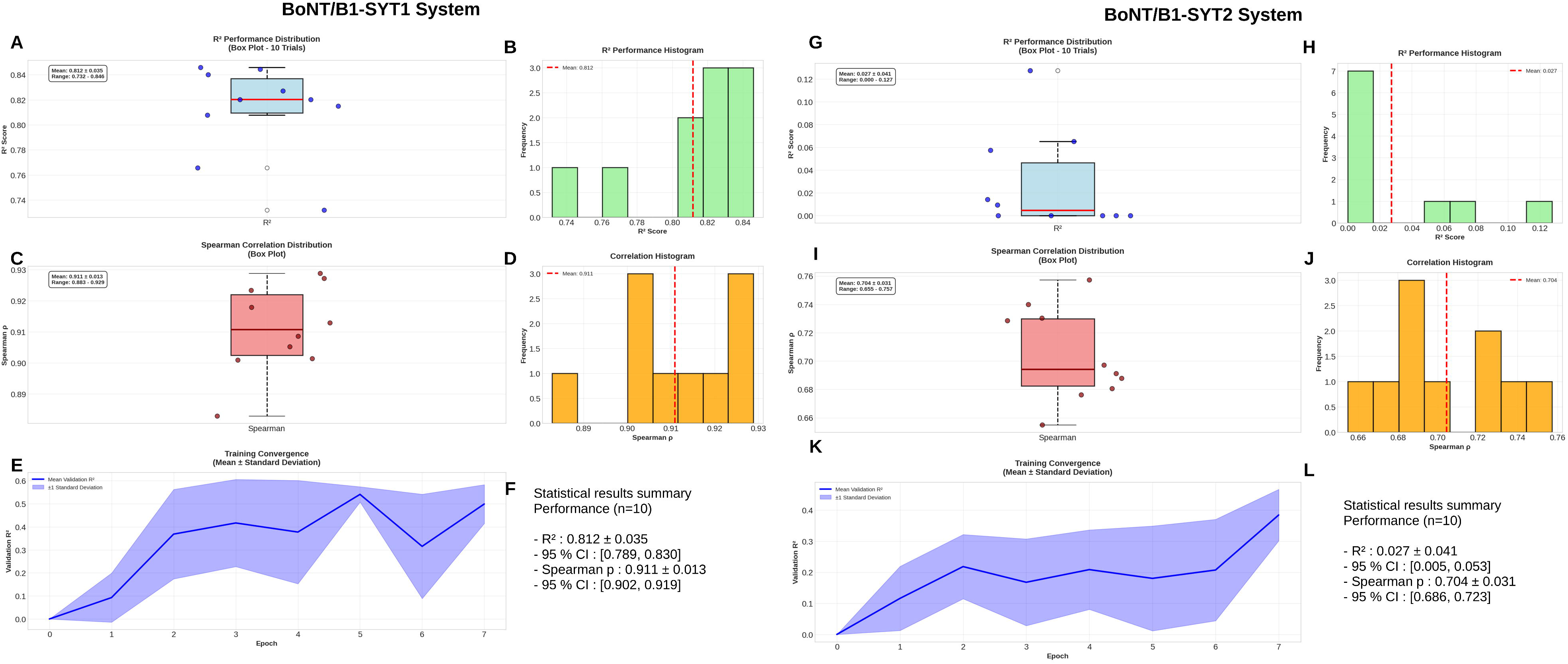

