## Supplementary material for "An AI-Driven Platform for Deconstructing and Engineering Biomolecular Recognition": Materiel and method, Figure S1-2, Table S1-2

**The PDF file includes:**

Materials and Methods

Figures S1 to S2

Tables S1 to S2

References

Materials and Methods

**Graph Representation of Biomolecular Structures.**

Feature engineering. We converted molecular dynamics simulation frames into graph neural network-compatible representations using a 13-dimensional feature vector per atom. 1) Electrostatic features (partial charges from force field topology (Charmm36m(1)), 2) mass properties (normalized atomic masses), 3) element encoding (atomic numbers: H=1, C=6, N=7, O=8, S=16 and P=15), 4) residue hydrophobicity (Kyte-Doolittle scale(2), normalized to [-1, 1]), 5) charge classification (positive=1, negative=-1, neutral=0), (6) structural indicators (backbone vs sidechain), 7) segment encoding (one-hot for biomolecule component referred as segid).

Edge construction. Two edge types were established:

Covalent edges: Directly from PSF topology files.

Non-covalent edges: Spatial proximity edges (< 6 Å) identified using k-d tree algorithms.

The edge features included:

. Radial basis functions (16 dimensions for distance encoding)

. Bond type indicators (covalent vs non-covalent: note that only inter-molecular non-covalent bonds are accepted, the intra-molecular are rejected (not necessary for our approach))

. Directional vectors (normalized atomic displacements)

**Neural Network Architecture**

We designed a graph convolutional network with frame interleaving. As shown in figure 1, the architecture includes 3 GCN layers (128→64→32 neurons) with batch normalization, 30% dropout for regularization, multi-head attention for interface-specific feature pooling and interface predictor linear layer for stability scoring. Interface detection was implemented using k-d tree spatial algorithms with a 6.0 Å threshold to identify contacting atoms between molecular segments.

**Training protocol**

We employed frame interleaving to ensure each data subset spanned the entire simulation trajectory: training frames (0, 3, 6...), validation frames (1, 4, 7...) and test frames (2, 5, 8...), each representing 33% of the data. Model optimization used AdamW (learning rate=0.001, weight decay=1e-4) with mean squared error loss and early stopping (patience=15 epochs). Statistical robustness was assessed through 10 independent training trials with different random seeds to evaluate model variance.

**Interface Perturbation Analysis**

We implemented six distinct biological perturbation strategies to assess interface criticality: 1) electrostatic charge reversal of Arg/Lys/Asp/Glu residues, 2) hydrophobicity inversion of hydrophobic core, 3) introduction of bulky sidechain substitutions, 4) disruption of π-stacking networks, 5) breaking of hydrogen bonding networks, and 6) structural displacement of interface atoms. Interface importance was quantified as Δ = |P(perturbed) - P(original)|, where P represents the model-predicted interface stability score. Residues were ranked by Sum Δ, which weights contributions by both interaction strength and frequency.

**Configurable Temporal Staging**

Temporal analysis in TPS is performed across user-defined stages. For clarity and consistency in this study, we devided trajectories into three equal-length regimes: early (initial binding events), mid (equilibrium interactions), and late (stabilized complex formation). However, TPS is designed with flexible stage definition: users may specify any number of stages, uneven time intervals and employ unsupervised clustering to identify natural kinetic phases directly from trajectory data. This flexibility allows TPS to model a wide range of dynamic processes including stepwise complex assembly, induced-fit binding, and dissociation pathways, by aligning the analysis with the underlying biochemical timeline.

**Statistical Validation**

We performed multi-trial statistical validation with 10 independent training runs to assess model variance. Statistical significance was evaluated using bootstrap confidence intervals (CI) (95% CI) across all trials. Performance metrics included R² coefficient (predictive accuracy), Spearman correlation (rank correlation with true stability), mean absolute error (absolute prediction error), and within-tolerance metrics (percentage of predictions within 10/15/20% of true values). The final model was selected based on balanced criteria of validation loss and test R² performance to ensure both training stability and predictive power.

**TPS-Lite: Accessible Static Structure Analysis**

For researchers without access to molecular dynamics simulations, we provide TPS-Lite, a simplified implementation requiring only a PDB structure and a PSF topology file. TPS-Lite performs static interface perturbation analysis using the same physicochemical feature engineering as full TPS (charge, hydrophobicity, structural indicators), by replaces the trained neural network with heuristic importance scoring based on residue properties and contact counts.

Given a protein complex structure, TPS-Lite: 1) identifies interface residues through spatial proximity (default cutoff: 6.0 Å), 2) calculates importance scores combining electrostatic magnitude (weight: 0.5), hydrophobicity (0.3), and classifies interaction types (electrostatic, hydrophobic, aromatic, hydrogen-bond, or general) based on residue chemistry. While lacking the temporal resolution and causal precision of full TPS, TPS-Lite enables rapid interface mapping suitable for mutagenesis guidance, educational use, and initial screening. The complete TPS-Lite implementation is included in my github.

**Intelligent Interface Optimization Scanner (IIOS)**

The Intelligent Interface Optimization Scanner (IIOS) is a standalone computational tool designed to convert interface analysis into actionable mutation suggestions for protein engineering. It operates independently of the TPS framework and requires only molecular graph files as input, though it also supports direct loading of PDB and PSF structural giles through MDAnalysis(3) integration for enhanced accessibility.

Interface residues were identified using atom-level distance checking with a user-definable cutoff (default: 5.0 Å). The framework implements a flexible chain selection system that accepts multiple identification formats including segment IDs (segid PROA), chain identifiers (chain A), residue ranges (resid 30:60), and complex Boolean combinations (segid PROA and resid 30:60). For each molecular dynamics frame, atomic positions were extracted from graph representations and filtered by chain assignment. Residue-residue contacts were detected using a k-d tree algorithm, and partner frequencies were calculated as: Frequency = (Unique frames with any interaction) / (Total frames analyzed).

Residues with partner frequencies below a configuration threshold (default: 0.1) were flagged as rare interactors and prioritized for mutation screening. The system also implements a proximity scanning mode that identifies “never-paired”residues within an extended distance cutoff (proximity_cutoff = interface_cutoff x proximity_factor, default: 1.5x), enabling the discovery of optimization opportunities beyond established interfaces.

IIOS employs a partner-aware suggestion that evaluates the chemical environment of each target residue through comprehensive neighbor analysis. For each candidate residue, the system calculates local chemical properties including charge distribution, aromatic character, polarity and hydrophobicity of all partners within the search radius. The system scans for four interaction types: 1) Salt-bridge optimization: Introduces complementary charges near oppositely charged partners (distance < 6.5 Å) with energy scoring based on distance-dependent electrostatic potentials. 2) Hydrogen-bond enhancement: Adds polar residues near hydrogen-bond donors or acceptors (distance < 4.0 Å) with geometry-appropriate substitutions (Ser, Thr, Asn, Gln, Tyr). 3) π-stacking interaction: Suggest aromatic substitution near existing aromatic partners (distance < 6.5 Å) with preference for Phe, Tyr, Trp, and His mutations. 4) Hydrophobic-core strengthening: Recommends aliphatic (Leu, Ile, Val, Met) or aromatic mutations in hydrophobic clusters to enhance non-polar packing interactions.

Each suggestion is assigned an expected ∆∆G value based on physicochemical propensity scores derived from empirical interaction energies, with penalties for steric clashes and backbone conformational strain. Confidence scores incorporate multiple factors including expected energy improvement magnitude (∆∆G), partner proximity, chemical complementarity, and structural accessibility. The framework supports single-point mutations as well as cooperative multi-residue optimization through configurable combination modes.

All user-adjustable parameters are centralized in a single “CONFIG” dictionary at the beginning of the code, including: File paths (graph, PDB, PSF), chain selection (target and partner with flexible identification syntax), mutation control (target chain, partner chain or both), residue exclusion criteria (aromatic preservation, alanine conservation), partner filtering requirements, partner analysis limits, results display options (target partners, or both), analysis parameters (frequency threshold, frame limits), output parameters (top N suggestions, export formats), ML model integration options, and mutation mode (single, double or triple mutation points). The modular architecture enables straightforward customization for specific protein engineering applications while maintaining consistent algorithmic behavior across diverse molecular systems.

**Code Implementation**

All analyses were implemented in Python 3.9 using PyTorch Geometric 2.3.0 for graph neural networks^10^, MDAnalysis 2.4.0 for trajectory processing, and scipy 1.10.0 for spatial algorithms. Visualization utilized matplotlib 3.7.0 and seaborn 0.12.0.

**Code Availability**

The full implementation of Temporal Perturbation Scanning (TPS), TPS-Lite, and Intelligent Interface Optimization Scanner (IIOS) including scripts for graph construction, neural network training, perturbation analysis, and visualization, is publicly available on GitHub: <https://github.com/fodil13>. All codes are released under a custom license permitting free academic use; commercial use requires a license.

**Figure S1**

**Fig 2.** The Perturbation Scanning (PS) Framework. (Upper pannel) Worflow for AI-driven deconstruction of biomolecular recognition. Molecular dynamics trajectories of a biomolecular complex (e.g, a neurotoxin bound (BoNT/B1 1150-1280, colored in white) to its receptor (synaptotagmin 1 (SYT1 30-85) or 2 (SYT2 30-85), colored in blue) are converted into a series of molecular graphs. Our graph creation code engineers 13-dimensional node features (including electrostatic, hydrophobic and steric properties) and constructs both covalent and non-covalent edges. (Lower panel) **Architecture:** The model processes the molecular graphs using a multi-layer Graph Convolutional Network (GCN). The architecture performs message passing along both covalent and non-covalent edges to learn a complex, hierarchical representation of the molecular system, culminating in a predicted stability score. **Frame Interleaving Training Strategy:** To respect the temporal correlation of MDS data, frames are split into training, validation and test sets using a consistent interleaving pattern (every 3^rd^ frame). This strategy prevents data leakage by avoiding contiguous blocks of time in a single split and crucially, forces the model to learn from and generalize across the entire ensemble of conformational states sampled by the simulation, rather than memorizing a specific temporal segment. **Multi-run validation:** The training process is repeated over multiple independent trials (n=10). Performance metrics (R², Spearman correlation) are aggregated, with results reported as mean ± standard deviation and confidence intervals, quantifying the pure variance of the model and guaranteeing the reliability of the findings.

**Figure S2**

**Fig 3.** Statistical performance comparison of the PS stability predictor for the BoNT/B1-SYT1 and BoNT/B1-SYT2 systems. For both systems, MD trajectories were converted into molecular graphs and processed using the same Temporal Interleaving training protocol. Each model was trained over 10 independent trials, and performance metrics were aggregated to quantify model variance. (A, G) Boxplots showing the distribution of test-set R² scores across 10 independent runs for SYT1 (A) and SYT2 (G). Individual data points represent each trial. (B, H) Histograms of the R² distributions for SYT1 (B) and SYT2 (H), with dashed lines marking the mean performance. (C, I) Boxplots summarizing Spearman rank correlation coefficients (ρ) across trials for SYT1 (C) and SYT2 (I). (D, J) Histograms of Spearman correlations for SYT1 (D) and SYT2 (J). (E, K) Training convergence curves showing the mean validation R² ± standard deviation across trials for SYT1 (E) and SYT2 (K). These curves reflect the stability and reproducibility of the optimization process. (F, L) Statistical summary panels reporting the means ± standard deviation and the 95% confidence intervals for R² and Sperman ρ for SYT1 (F) and SYT2 (L) systems. Together, these panels compare how the PS learns stability patterns for each receptor. The same protocol was used for SYT1 and SYT2; differences in model performance arise solely from differences in the underlying MD conformational ensembles.

Table S1.

Total PS ∆ values for plasmolipin in different membrane environments

| System | Early Sum ∆ | Mid Sum ∆ | Late Sum ∆ |
| --- | --- | --- | --- |
| Plasmolipin-Lipid Raft | 33,987 | 32,03 | 31,753 |
| Plasmolipin-POPC | 8,37 | 8,657 | 8,465 |

Table S2.

Lipid composition within 6 Å of plasmolipin and corresponding TPS Δ interaction scores in the lipid raft system. Percentages indicate relative abundance of each lipid type within the contact shell.

| Lipid | Contact Frequency (%) | TPS Δ (sum) | Δ per Contact |
| --- | --- | --- | --- |
| POPC | 44.3 | 10.06 | 0.047 |
| CER160 | 13.1 | 16.48 | 0.262 |
| CHL1 | 15.4 | 6.07 | 0.082 |
| BGLC | 9.8 | 7.74 | 0.165 |
| BGAL | 5.8 | 3.25 | 0.116 |
| ANE5AC | 5.0 | 1.43 | 0.060 |
| BGALNA | 4.0 | 1.34 | 0.071 |
| BGAL | 2.7 | 0.65 | 0.050 |
